# The neural signature of numerosity: Separating numerical and continuous magnitude extraction in visual cortex with frequency-tagged EEG

**DOI:** 10.1101/802744

**Authors:** Amandine Van Rinsveld, Mathieu Guillaume, Peter J. Kohler, Christine Schiltz, Wim Gevers, Alain Content

## Abstract

The ability to handle approximate quantities, or *number sense*, has been recurrently linked to mathematical skills, though the nature of the mechanism allowing to extract numerical information (i.e., numerosity) from environmental stimuli is still debated. A set of objects is indeed not only characterized by its numerosity but also by other features, such as the summed area occupied by the elements, which often covary with numerosity. These intrinsic relations between numerosity and non-numerical magnitudes led some authors to argue that numerosity is not independently processed but extracted through a weighting of continuous magnitudes. This view cannot be properly tested through classic behavioral and neuroimaging approaches due to these intrinsic correlations. The current study used a frequency-tagging EEG approach to separately measure responses to numerosity as well as to continuous magnitudes. We recorded occipital responses to numerosity, total area, and convex hull changes but not to density and dot size. We additionally applied a model predicting primary visual cortex responses to the set of stimuli. The model output was closely aligned with our electrophysiological data, since it predicted discrimination only for numerosity, total area, and convex hull. Our findings thus demonstrate that numerosity can be independently processed at an early stage in the visual cortex, even when completely isolated from other magnitude changes. The similar implicit discrimination for numerosity as for some continuous magnitudes, which correspond to basic visual percepts, shows that both can be extracted independently, hence substantiating the nature of numerosity as a primary feature of the visual scene.

## Introduction

When it comes to sets of more than five objects, we can rapidly figure out an approximation of the number of items, the *numerosity*, without counting [1]. Humans share with other animal species an *intuition* for numerical quantities [2]. The nature of the cognitive mechanism behind this ability to approximate large numerosities is still vividly debated. Researchers largely assume that we possess an *Approximate Number System* (ANS), a specific system that extracts numerosity and builds a mental representation of the discrete numerical magnitude from the visual scene [3]. However, a set of objects is characterized not only by numerosity, but also by several *continuous* visual features: the individual object sizes, the extent of the set, etc. These continuous magnitude dimensions are intrinsically related to numerosity (e.g., a more numerous set naturally occupies a larger area), and may serve as critical visual cues to access numerosity. This has led some authors to suggest that there is no specific cognitive mechanism devoted to number processing and that numerosity is either processed by general magnitude mechanisms or emerges from a combination of continuous dimensions [4]. No consensus has been reached thus far on how continuous magnitudes contribute to numerosity processing, and a large body of evidence has demonstrated that they can either facilitate or interfere with numerosity judgments [5]. The current study capitalized on a frequency-tagging electrophysiological approach to isolate numerosity from continuous magnitude dimensions and to measure the specific cerebral responses driven by both.

The ability to discriminate sets of objects based on numerosity is thought to be shared with other animal species and to be present in infants long before the development of language [6 7 8]. There is substantial behavioral and neuroimaging evidence of this numerical ability. For instance, recent experiments highlighted a spontaneous bias in favor of numerosity against continuous magnitudes when participants had to choose the odd one out of three collections of dots or to categorize collections as “large” or “small” [9 10]: in both cases numerosity was spontaneously selected as the decision criterion. Further, some studies identified populations of neurons in the parietal cortex of humans and monkeys that are specifically tuned to numerosity [11 12]. Nevertheless, the exact nature of the mechanism supporting this numerical ability remains largely unknown, due to the inherent correlation of continuous magnitude changes with numerosity changes. Continuous magnitude rather than numerosity itself could account for the observed results [13]. It is an open question whether the cognitive system is able to rapidly extract numerosity information necessary to build up a representation independent of continuous magnitude variations – and if the system has that ability, what happens to the co-varying continuous magnitude information as numerosity is processed? The ANS theory proposes that all continuous magnitudes are filtered out during a *normalization* stage [2], but there is not much evidence about this filtering stage since continuous magnitudes substantially affect numerosity judgments [14].

Alternative theories have proposed that numerosity is yoked to continuous magnitude processing. Among these, a Theory of Magnitude (ATOM) describes the relationship between continuous magnitudes and numerosity in terms of a unique system that is capable of representing any kind of discrete and continuous magnitude, including numerosity, time (duration) and space (extension) [15]. Some authors proposed a general *sense of magnitude* for both continuous and discrete quantities, in which size perception is developmentally and evolutionarily more primitive than numerosity, and continuous magnitude plays a key role in the development of numerical magnitude processing [16 17]. There is substantial empirical evidence supporting both common and separate neural areas for numerical and continuous magnitudes (e.g., [18 19 20 21]). Partially overlapping topographical maps for numerosity and continuous magnitude extraction were identified within the human parietal cortex, although distinct neural tuning and organization within the maps suggested distinct processing mechanisms [22 23]. Within these overlapping areas, the right parietal lobule was identified as a likely anatomical location of the generalized magnitude processing system, according to a recent fMRI meta-analysis [24]. Further, some authors argued that numerosity is only an abstract cognitive construct resulting from the weighting of all continuous magnitude features present in the visual stimulus, and that numerosity is extracted through adaptive recombination of lower-level sensory information according to the needs of a particular context [4]. Such *Sensory Integration* (SI) theory assumes that all existing evidence of numerosity extraction can be explained by cognitive control mechanisms handling the integration of continuous magnitudes.

The major challenge in disentangling these hypotheses and in understanding the mechanism of numerosity processing is to isolate numerosity from continuous magnitudes. Several elegant ways to control for continuous dimensions have been developed for behavioral tasks [25 26 27 28], but they control for all magnitude changes across the stimulus set, though each stimulus still contains information about both numerosity and continuous dimensions. Indeed, any visual stimulus carries information about both numerosity and continuous magnitudes. These methods are thus incapable of disentangling numerosity from non-numerical magnitude processing in any strict sense. Importantly, this limitation applies to almost – if not all – evidence provided so far in favor of the ANS theory.

The current study used a frequency-tagging approach which consists in recording Steady-State Visual Evoked Potentials (SSVEP) corresponding to the neural responses that are specific to periodic stimulus changes in a single given dimension [29]. SSVEP have already been successfully recorded in response to numerosity variations [30] but the present study is the first to systematically isolate the discrimination of numerosity and of continuous magnitudes with a novel frequency-tagged experimental paradigm that requires no explicit task (and thus no decision nor judgment): Visual stimulation followed an oddball paradigm in which a deviant stimulus was periodically introduced within a stream of standard stimuli [31]. Critically, we strictly manipulated the nature of the periodic change, so that only the dimension under consideration periodically fluctuated [32]. This manipulation allows for the recording of the neural responses synchronized to changes in the target dimension, since only that particular dimension was updated periodically. The current design allows to track neural discrimination of changes in numerosity as well as in each of the continuous dimensions, by assigning each dimension to be the periodic deviant in separate experimental conditions. If the visual system discriminates periodic changes relative to the fluctuating dimension, the brain should produce responses synchronized to the deviant frequency and its harmonics [33]. We are thus able to record brain activity specifically related to the discrimination of numerosity and of each continuous dimension.

## Experimental design

Sequences of dot arrays were presented, updating at a base rate of 10 Hz (i.e., 10 dot arrays per second). Each presented stimulus was characterized by five dimensions: number of dots, dot size, total occupied area, convex hull (i.e., the surface of the smallest set enclosing all dots of the collection), and density. The dot arrays were made to vary randomly along all dimensions except one, which changed systematically at a rate of 1.25 Hz (i.e., one deviation every eight items). On each 44-second block, the periodic dimension was made to be either numerosity or one out of the four continuous magnitudes (i.e., dot size, total area, convex hull and density), see Figure 1. The ratio by which the deviant dimension varied from the standards could be 1.1, 1.2, 1.3, 1.4 or 1.5, leading to a design consisting of five dimensions and five ratios. Each of the 25 conditions was repeated in 3 blocks. As the current electrophysiological recordings did not allow to precisely identify where the discrimination occurs along the occipito-parietal stream, we further assessed whether this discrimination could theoretically be supported by early visual regions. Therefore, we ran our stimuli through a relatively simple model of early visual processing to assess whether the model would predict the discrimination we observed in the electrophysiological results.

**Figure 1.**
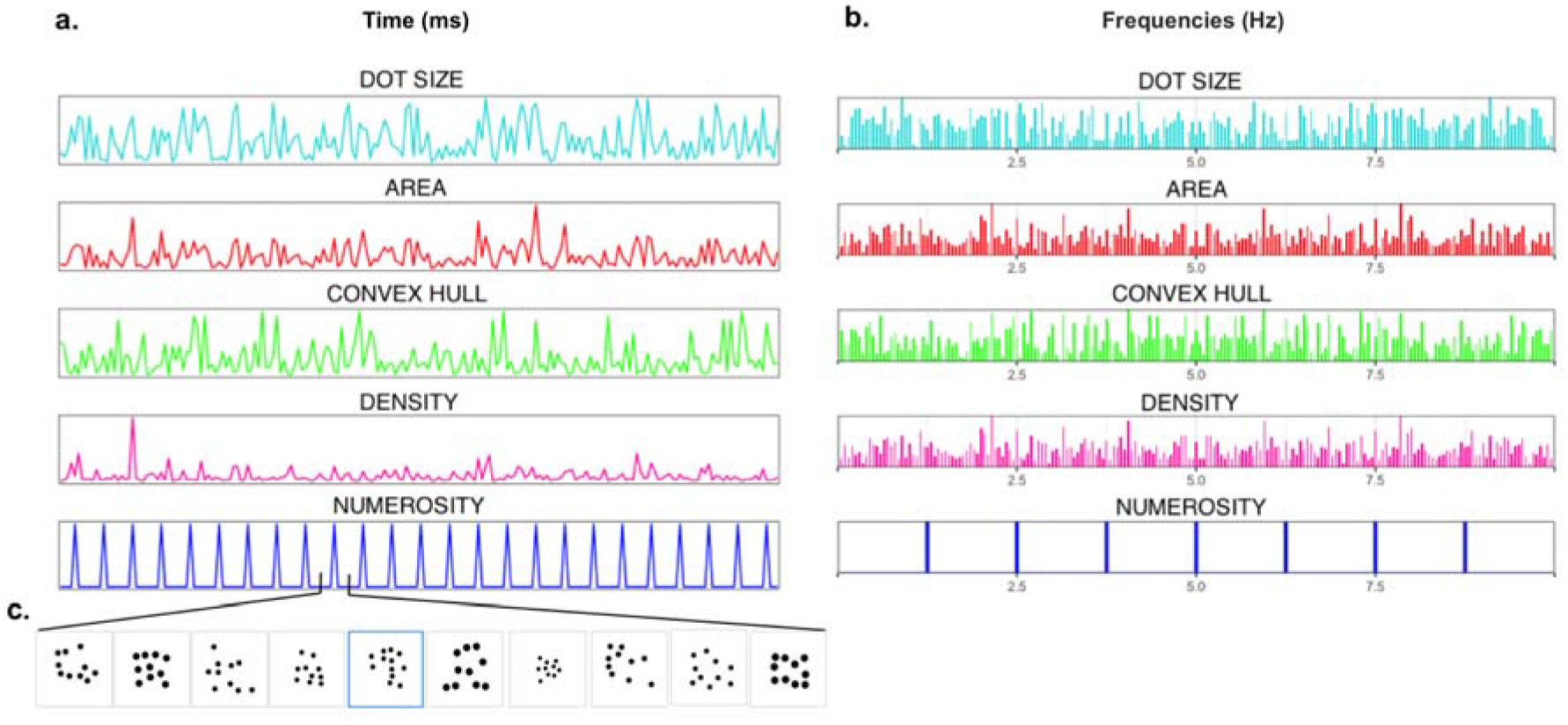
Illustration of the experimental design, example with numerosity as the periodic deviant: **a**. Sequences of dot patterns were characterized by five dimensions: dot size, total area, convex hull, density, and numerosity. Time series of values for the five dimensions during the first 20 seconds of a trial (200 stimuli) are plotted. In this example, numerosity is the deviant that varies periodically at 1.25 Hz while the continuous magnitude dimensions vary randomly. **b**. Frequency spectra of the corresponding time series after Fourier transformation. Only the periodic varying dimension, in this example numerosity, produces clear spectral power at 1.25 Hz and higher harmonics. **c**. Illustration of a subset of the stimuli.

## Results

To measure the neural responses corresponding to the discrimination of each dimension, we summed the baseline-corrected amplitudes of the target frequency (1.25 Hz) and its harmonics up to the eighteenth (i.e., the highest harmonic with a significant response, see Methods), excluding harmonics of the base rate frequency (i.e., 10 and 20 Hz, as in previous studies [34]). The Sums of Baseline-corrected Amplitudes (SBA) were computed per participant and per condition, and then averaged at the group level. We found a clear response to the deviant stimulus at the group level for the largest ratio of three visual dimensions: Total Area, Convex Hull, and Numerosity (Figure 2). The strongest SBA peaks were recorded around medial occipital electrode Oz, which is consistent with previous results on number discrimination [32]. The mean values of the SBA for the highest ratio (1.5) averaged across the whole posterior scalp and their corresponding 95% confidence intervals were respectively 0.27 μv [0.23, 0.31] for Total Area, 0.25 μv [0.21, 0.28] for Numerosity, 0.23 μv [0.19, 0.27] for Convex Hull, 0.12 μv [0.08, 0.15] for Density, and 0.02 μv [0.00, 0.05] for Item Size. To get a clearer picture of these results, we considered four posterior regions of interest for further analyses: the medial occipital, medial occipito-parietal, left occipito-parietal and right occipito-parietal regions (see Methods).

**Figure 2.**
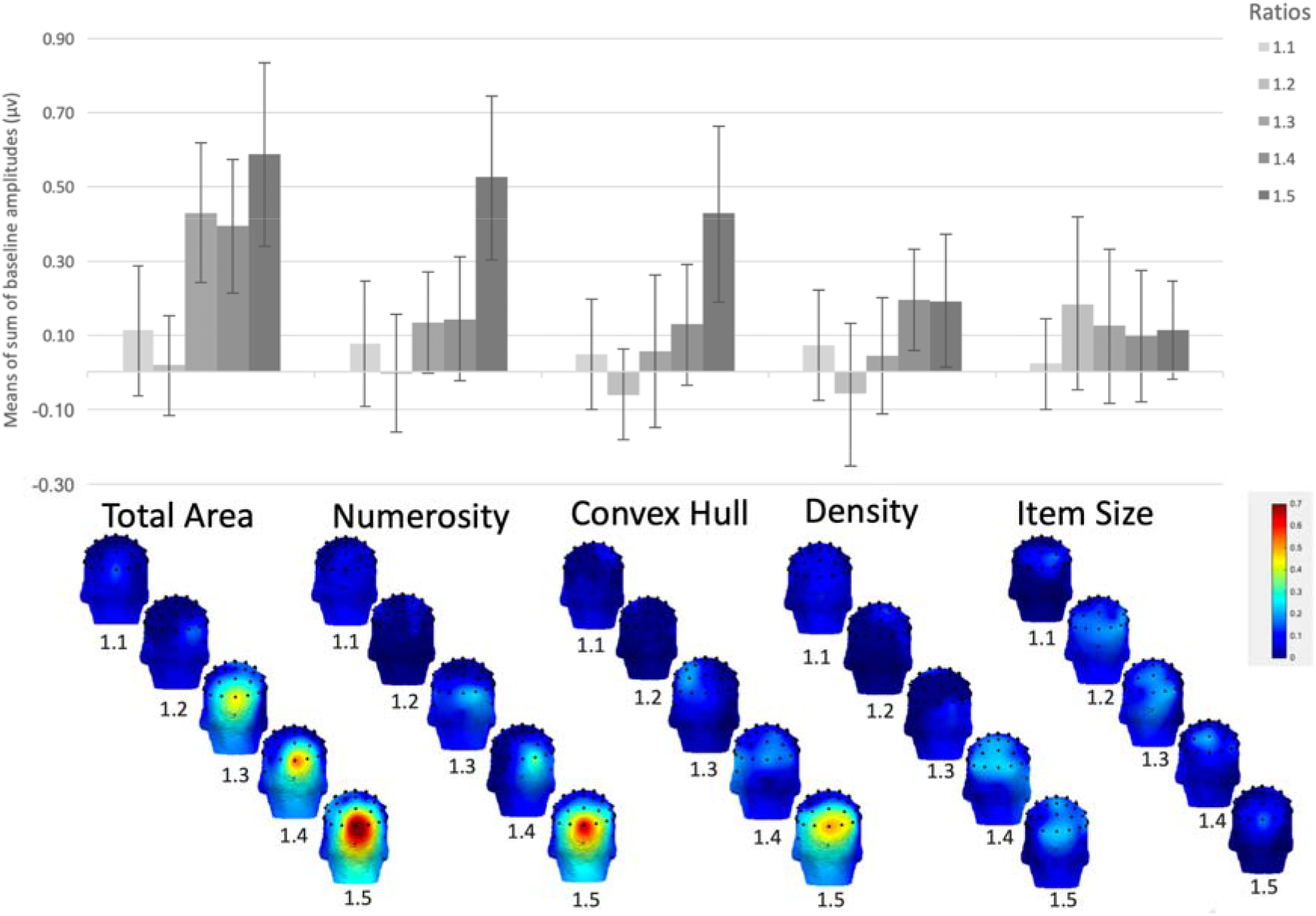
Upper panel: Average of the sums of the baseline-corrected amplitudes (SBA, in microvolts) for every condition, as a function of the change ratio. The SBA is the sum of the target frequency (1.25Hz) and its harmonics, excluding the base rate (see text). Error bars depict 95% Confidence Intervals. Lower panel: Scalp topographies of the SBAs (in microvolts) for every condition, as a function of the ratio.

To evaluate the effect of the ratio manipulation, we constructed a linear mixed model, with the ratio and the regions as fixed predictors of the SBA, and participants as random factor. Visual inspection of residual plots did not reveal any obvious deviations from homoscedasticity or normality. We compared the full model to two reduced models that did not include the ratio and region predictor, respectively, using chi-square tests on the log-likelihood values. For Numerosity, the full model had a better fit than both reduced models, χ^2^(1) = 21.60, *p* < .001, and χ^2^(3) = 10.874, *p* = .01, respectively without ratio and without regions. A similar result was observed for Total Area where the full model had a better fit than both reduced models, χ^2^(1) = 23.915, *p* < .001, and χ^2^(3) = 31.077, *p* < .001,respectively without ratio and without regions. We thus observed a significant ratio effect in both conditions. The strongest responses to periodic changes in Numerosity and in Total Area were recorded in the medial occipital electrodes. For Convex Hull, the full model had a better fit than the reduced models without ratio, χ^2^(1) = 35.53, *p* < .001, but not better than the reduced model without regions, χ^2^(3) = 2.66, *p* = .45. For Density, the full model had a barely significantly better fit than the model without ratio, χ^2^(1) = 4.07, *p* = .04, but not than the one without regions, χ^2^(3) = 0.171, *p* > .10. Finally, for Size, the full model did not have a significantly better fit than either reduced models, χ^2^(1) = 0.171, *p* > .10, and χ^2^(3) = 4.55, *p* = .21.

The linear mixed models thus showed that the brain response to the periodic change of Numerosity and Total Area was significantly affected by both the ratio between deviant and standard and by the location of the electrodes (see Figure 2). The effect was largely driven by responses in the medial occipital region: At the group level, the medial occipital region produced a clear SBA response to periodic changes in Area when the ratio was 1.3 or more (mean SBA for ratio 1.3 = 0.43 μv; 95% CI [0.24, 0.62]). Periodic changes in Numerosity also produced clear SBA responses in the medial occipital region, although only for the largest ratio of 1.5 (mean SBA = 0.53 μv; 95% CI [0.30, 0.74]). Convex Hull and density both yielded significant effects of ratio, but not of region, but in both conditions the medial occipital region produced the strongest responses. For Convex Hull, only ratio 1.5 reached significance (mean SBA = 0.43 μv; 95% CI [0.19, 0.61]), while ratio 1.4 of Density (mean SBA = 0.19 μv (95% CI [0.05, 0.33]) was just at the limit of significance. However, there was no significant brain response to periodic changes of Item Size (mean SBA = 0.11 μv (95% CI [−0.02, 0.25]), consistent with the lack of region or ratio effect in the linear mixed effect model.

To enrich our interpretation of the electrophysiological measurements, we ran an additional analysis that was aimed at testing whether a relatively simple model of early visual processing, which was not designed to account for number processing, would predict the results we observed. To that end, we applied a second-order contrast (SOC) model (see Methods) to the images used in the experiments. The model output was closely aligned with our SSVEP data: the standard and deviant images produce systematically different predicted responses for Numerosity, Total Area and Convex Hull, the three dimensions that elicited synchronized brain responses, but not for Dot Size and Density, the two dimensions that did not, see Figure 3 (see Table 1 of supplementary material for detailed results). It is important to note that the model was designed to capture activity elicited in the visual cortex by simple gratings, textures and natural scenes, and fit to functional MRI data, a different neuroimaging modality from the one we used. The fact that the output of such a model is so well-aligned with our measured data is good news both for the model and for our approach. The model’s ability to approximate our measured electrophysiological responses to numerosity, despite not being designed to capture or account for numerosity, is especially compelling, and may indicate that the initial stages of numerosity processing rely on relatively simple computations taking place in early visual cortex.

**Figure 3.**
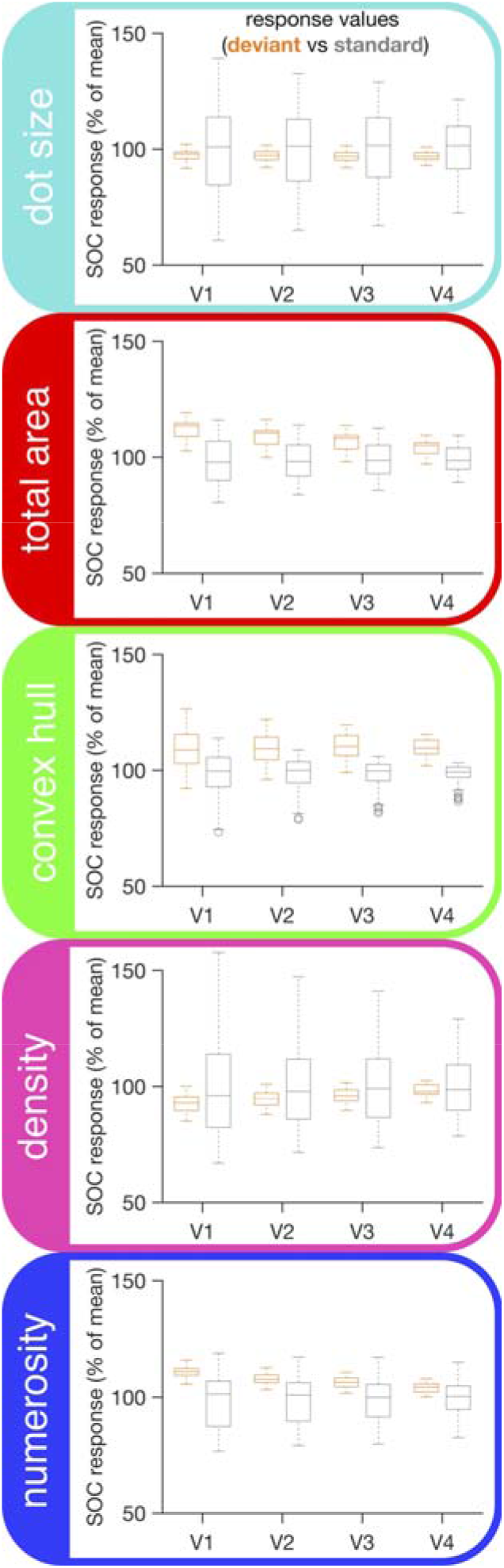
For each periodic deviant indicated with the box label on the left, the graphs depict the average responses in four visual areas, as predicted by the SOC model. Each box represents a separate 1.5 deviant to standard ratio. For each boxplot, the central mark is the median, the edges of the box are the 25th and 75th percentiles, the whiskers extend to the most extreme data points not considered outliers, and the outliers are indicated with dots.

## Discussion

The objective of the current study was to isolate and contrast the specific responses to changes of numerosity and of four continuous visual magnitudes. We observed clear electrophysiological responses that were synchronized to the frequency of a periodically occurring deviant stimulus, when that deviant encompassed a change of numerosity. These synchronized responses support the view that the human brain can spontaneously discriminate periodic changes in numerosity. The observed effect of region of interest and the results from the model approach further suggest that this ability is primarily driven by occipital cortex. The frequency-tagging approach taken in the current study allows a de-correlation of numerosity and continuous magnitudes, which means that we could choose natural dimensions as a strong comparison point in terms of low-level changes in visual features. Indeed, we demonstrated that when numerosity was totally de-correlated from continuous magnitudes at the sequence level, its changes could still be discriminated as automatically as changes in Total Area and Convex Hull.

These findings are in line with recent studies that have identified rapid electrophysiological responses to numerosity (75 ms after stimulus presentation) [35] and with evidence that encoding of numerosity occurs very early in the visual processing stream: tuning for numerosity has been demonstrated in early visual regions of occipital cortex (V1, V2, and V3) [30 32 36 37]. Crucially, as our experimental design ensured that only a single dimension varied periodically over a given stimulus sequence, the current observed responses can be uniquely associated with numerosity, and contributions from any of the continuous dimensions in isolation can be ruled out. It is worth noting that our study involved passive viewing only. Hence, our findings do not preclude the existence of later interactions between the dimensions, which have been recurrently reported behaviorally in non-symbolic comparison tasks involving decision-making and cognitive control. It is thus still possible that a weighting [4] or normalization of the various magnitude information [2] occurs at later processing stages while performing a numerosity task (e.g., non-symbolic estimation or comparison tasks). Recent studies support the view that such interactions are deliberate, and strategic, rather than perceptual [38]. In other words, the current results do not support the hypothesis that the number of items in a visual scene is initially processed through an active weighting of continuous magnitudes at that early processing stage.

The results of the conditions in which continuous magnitudes were manipulated, revealed synchronized responses that could be uniquely associated with the total area occupied by the dots and the extent of the convex hull. On the contrary, no such synchronizations were observed for periodic variations of density and of dot size. This pattern of results indicates that the brain can spontaneously discriminate total area and convex hull, but not density and dot size: Total Area and Convex Hull are both directly related to low-level visual features: Total Area is confounded with stimulus luminance, a primary property of the visual scene [39] to which the visual system responds in a nearly veridical manner [40]. Convex Hull is confounded with the size of the space taken in the visual scene, and thus with the width and height of the visual angle sustained by the entire stimulus, properties that are known to influence the brain response to the stimulation [41 42]. Human vision thus seems to be provided with an early discrimination mechanism for numerosity that operates in an equivalent way to the mechanisms involved with decoding low-level visual features, suggesting that numerosity may also be considered a primary visual feature [43]. The topological invariance of numerosity has been proposed as a key visual attribute distinguishing numerosity from continuous magnitudes [44 45]. This unique aspect of numerosity might be key to its utility as a primary feature of visual scenes.

The SOC model was designed to capture activity elicited in visual cortex by relatively simple image stimuli, and fit to functional MRI data. The responses predicted by the model are well-aligned with our electrophysiological data, so that variability over dimensions that produced synchronized brain responses also produced systematically different predicted model responses. This indicates that the synchronized brain responses could plausibly be elicited by the mechanisms in early visual cortex that the model was designed to capture. It is perhaps not surprising that early visual cortex would be sensitive to Total Area and Convex Hull, as indicated by the model. However, stimuli that are made to vary along the numerosity, but not the other dimensions, also elicit systematically different predicted model responses. This finding is more remarkable as the model was not designed to detect that kind of change. It thus offers support for the idea that numerosity is a primary visual feature that, at least at initial processing stages, relies on relatively simple computations taking place in early visual cortex.

It is worth noting that the observed discrepancies across dimensions suggests that the observed synchronized electrophysiological responses do not reflect a general mechanism of response to any periodic deviant, but rather depend critically on the brain’s sensitivity to the change [33]. The results indicate that the brain is sensitive to Numerosity, Total Area and Convex Hull, while we found no evidence of sensitivity to periodic changes in Dot Size or Density. The lack of a density effect might be due to the range of number of dots used in the present experiment. Indeed, some authors have argued that the density of a dot array becomes salient only when the number of dots is much larger than the range we used (i.e., over hundreds of dots) [46]. According to their results, electrophysiological responses to density changes would not occur with arrays with less than fifty dots, which is consistent with our observations. The lack of a dot size effect could be explained by attention being allocated to the global visual scene rather than to its individual parts. The visual system can segment a scene to extract relevant information and previous research suggests that there would be no encoding of individual dots but rather an efficient general description of the scene [43]. Thus, although we did not observe specific responses to variations in dot size and density, these properties may be measurable under different experimental settings, including larger ratios or longer presentation times, with the latter manipulation possibly allowing for directed attention towards individual scene parts.

Taken together, the current results provide evidence in favour of the Visual Number Sense idea proposed by Burr and Ross, which considers numerosity a primary visual property that can be extracted from the visual scene [47]. Further support comes from recent evidence highlighting a specific neural sensitivity to numerosity very early in the visual stream, which was interpreted as direct and automatic encoding of numerosity information [48 49]. The seminal ANS theory proposed that lower-level visual features related to continuous magnitudes must be neutralized in a so-called normalization stage which precedes the extraction of number in an abstract, modality-free manner (i.e., the “abstract numerosity” as described by Gebuis [4]). Recent findings of neural sensitivity to numerosity both in early and late ERP components were interpreted as neural evidence of this normalization stage occurring in primary visual cortex prior to later summation stage [30]. The data presented here can also be considered as evidence of an early summation stage in primary visual cortex [37]. The current results demonstrate for the first time that a specific discrimination based on numerosity and some continuous magnitudes is possible similarly and very early in the visual stream. It is important to note that in a natural context, where the dimensions are not isolated as in the current experimental paradigm, and thus strongly correlate, early stage visual discrimination would likely be much stronger.

### Conclusion

The current study reports isolated measures of the brain’s ability to detect changes in both numerosity and continuous magnitudes, without the confounds that usually arise due to the correlation between these dimensions. The results show that numerosity can be rapidly discriminated in the visual stream independently of other visual features, supporting the hypothesis of an early visual number sense. We further suggest that numerosity is a primary attribute that can be directly extracted from the visual scene. Future research is needed to determine whether the ability to extract numerosity directly is an innate ability, or is learned over the course of visual development.

## Methods

### Participants

Twenty-five undergraduate students participated in the study. Volunteers suffering from or with a history of suffering from any neurological or neuropsychological disease, from any learning disability such as dyscalculia, or from any uncorrected visual impairment were not allowed to participate. We excluded four participants due to the presence of substantial noise in their EEG signal (*e.g*., noise due to transpiration or movements). The final sample thus consisted of twenty-one participants, with a mean age of 23.5 years (Standard Deviation, *SD* = 2.7, 9 females). Due to the length of the experiment, some participants failed to response to the colour change task (see below) and were excluded for some of the conditions, resulting in a final sample of 19 for Area, 18 for Convex hull, 18 for Numerosity, 19 for Size, and 20 for Density. No participant was excluded for more than two conditions. We followed APA ethical standards to conduct the present study. The Faculty Ethics Committee approved the methodology and the implementation of the experiment before the start of data collection. The experiment lasted two hours in total, and participants received 20 euros for their participation.

### Fast Periodic Visual Stimulation

#### Stimuli

To create the dot arrays, we used a preliminary version of the NASCO software [50]. NASCO is an open and freely usable MATLAB application that allows the generation of dot arrays while keeping a given dimension constant. Critically for the purpose of our study, for every condition, we generated 400 standard pictures in which only one dimension (determined by the condition under consideration) was kept constant while other features stochastically varied. We then created 100 deviant arrays for each condition with a similar approach, by using the ratio of the given condition to determine the value of the only manipulated fluctuating dimension. After the creation of the pictures, we generated multiple sequences of 440 of these (since the whole stimulation sequence had 10 pictures per second during 44 seconds), including 385 frequent and 55 deviant stimuli. In order to statistically verify that stochastic fluctuations related to irrelevant dimensions were not periodic within these sequences, we computed the Fast-Fourier Transform (FFT) of the values taken by each dimension over time. We transformed the spectra into Z-scores and we averaged the values of the deviant frequency rate and its harmonics up to the seventh (1.25, 2.50, 3.75, 5.00, 6.25, 7.50, 8.75 Hz). We then iteratively selected for each condition one sequence in which the averaged periodicity of the dimension of interest was larger than the 99% probability unilateral threshold of a standard normal distribution (i.e., 2.32), and in which the averaged periodicity of the other four dimensions were less than this threshold. The exact same sequence was used thrice within a given condition to avoid any periodic artefact when averaging the repetitions.

#### Apparatus

We used MATLAB (The MathWorks) with the Psychophysics Toolbox extensions [51 52] to display the stimuli and record behavioural data. The EEG recording took place in a shielded Faraday cage (275 cm × 195 cm × 280 cm). Participants were comfortably seated at 1 meter from the screen, with their gaze in front to the centre of the screen (24’’ LED monitor, 100 Hertz (Hz) refresh rate, 1 ms response time). Screen resolution was 1024 × 768 px. The order of the conditions during the EEG recording session was counterbalanced across participants.

#### Procedure

Participants were instructed to look at the entire screen by keeping their gaze on a small fixation diamond that was continuously displayed at the centre of the screen. The fixation randomly changed colour from blue to red four to six times during a sequence and participants were instructed to press a button with their right forefinger each time they detected the colour change. Their responses were recorded to quantify compliance with the instructions. Participants’ mean response rates to the color change of the fixation diamond was 96%. No participant missed the color change more than once during a 44 s block. Such high detection rate indicates that participants followed the instructions and kept their gaze on the center of the screen during EEG acquisition.

Stimuli subtended a maximal visual angle of nine degrees. Stimulus presentation followed a sinusoidal contrast modulation from 0% to 100% [53 54]. The base frequency rate was 10 Hz, so that ten stimuli were displayed per second (and consequently each stimulus lasted 100 ms in total from onset to total offset). Every stimulation sequence lasted 48 seconds, including 44 seconds of recording and 2 seconds of fade-in and fade-out, which were not analysed.

During each sequence, dot arrays entailed one specific feature that was kept constant, and a periodic deviation from this constant every eight items. In other words, *deviant* stimuli were periodically displayed within a stream of *standard* dot arrays at the frequency rate of 1.25 Hz (see Figure 1). We manipulated two factors: the nature of the dimension that periodically fluctuated, and the ratio of the fluctuation from the standard to the deviant arrays. As for the dimension, we manipulated the Number (N) of dots, the individual dot Size (S), the total Area (A) occupied by the dots, the area of the Convex Hull (CH), and finally the Density (D) of the array. As for the ratio, we created dot arrays that deviated from the standard with five different ratios (1.1, 1.2, 1.3, 1.4, and 1.5). The manipulated dimension and the deviation ratio were fixed during a block. Therefore, by the combination of both factors we obtained twenty-five different conditions. Each condition was repeated three times, for a total of seventy-five 44 s stimulation sequences. When debriefed at the end of the experiment, participants reported changes of the dot arrays but no participant noticed periodic changes in any dimension.

#### EEG recording

EEG data was acquired at 1024 Hz using a 64-channel BioSemi ActiveTwo system (BioSemi B. V., Amsterdam, The Netherlands). The electrodes were positioned on the cap according to the standard 10-20 system locations (for exact position coordinates, see http://www.biosemi.com). Two additional electrodes, the Common Mode Sense (CMS) active electrode and the Driven Right Leg (DRL) passive electrode, were respectively used as reference and ground electrodes. Offsets of the electrodes, referenced to the CMS, were held below 40 mV. Eye movements were monitored with four flat-type electrodes; two were placed above and below participant’s right eye, the other two were positioned lateral to the external canthi.

#### Data analyses

Analyses were conducted with the help of *Letswave 6* (http://nocions.webnode.com/letswave). Data files were down-sampled from 1024 Hz to 512 Hz for faster processing. Data were filtered with a 4-order band-pass Butterworth filter (0.1 to 100 Hz) and re-referenced to the common average.

The fade-in and fade-out periods were excluded from the analyses leading to the segmentation of an EEG signal of 44 seconds (corresponding to the display of 440 stimuli). The three repetitions were averaged per condition and per participant. A Fast Fourier Transform (FFT) was applied on the signal to extract amplitude spectra for the 64 channels with a frequency resolution (the size of the frequency bins) of 0.016 Hz.

Based on the frequency spectra, we computed two measures to determine whether and how the brain specifically responded to the deviant frequency in the five conditions: Sum of baseline-corrected amplitudes, and Z-scores. The Sum of Baseline-corrected Amplitudes is expressed in microvolts and can thus quantify changes within the EEG signal [32 34]. We computed the baseline-corrected amplitudes by subtracting from each bin of interest (*i.e*., 1.25 Hz and its harmonics up to the eighteenth) the mean amplitude of their respective forty-four surrounding bins (excluding the immediately adjacent ones, and the two most extreme values), and we summed these values. We grouped electrodes of interest in four posterior regions of interest for further analyses: the medial occipital (O1, O2, Oz, Iz), medial occipito-parietal (Pz, POz, P1, P2, P3, P4, PO3, PO4), left occipito-parietal (P5, P7, P9, PO7) and right occipito-parietal (P6, P8, P10, PO8) regions [55].

### Visual cortex response predictions

We applied a cascaded, feedforward model of BOLD responses to visual stimuli, the second-order contrast (SOC) model to the images used in the conditions with the largest ratio (1.5). This model first passed the image through a bank of contrast-normalized, localized, V1-like filters, and then reprocessed the output in a second stage that, among other things, measured the contrast variability in the output of the first stage [56]. The SOC model has been fit to data for V1, V2, V3, and V4, and is effective at predicting how these areas respond to simple visual stimuli, including gratings and textures, while also capturing increased sensitivity to the structure of natural scenes in extrastriate visual areas. These features make it a reasonable model for estimating the activity that our dot images would elicit in early visual cortex. We applied the SOC model separately to each image, using model parameters for V1, V2, V3, and V4 provided in the original publication. We then summed the model output across each image to get a predicted average response of each brain area to each image update. This allowed us to compare the predicted average responses to the standard and deviant image updates, and use the comparison as an estimate of how strongly the deviants would be expected to drive synchronized brain responses in early visual cortex, according to the SOC model.

## Acknowledgement

This research was funded by the European Union’s Horizon 2020 research and innovation program under the Marie Skłodowska-Curie grant agreement No 799171 to AV and by PDR project No T.1052.15 and equipment credit No 2.5020.12 of Fonds National de la Recherche Scientifique to AC and WG. The authors declare no conflict of interest that might be interpreted as influencing the research, and APA ethical standards were followed in the conduct of this work.

