## Supplementary information for "The neural signature of numerosity: Separating numerical and continuous magnitude extraction in visual cortex with frequency-tagged EEG"

**Results of the Base Rate Responses (10 Hz)**

We recorded large responses at the base stimulation rate (10 Hz) at posterior electrode sites. We computed the Z-transformation of the amplitude of the base stimulation rate relative to its surrounding twenty bins (ten on each side) for every condition. Z values above the threshold of 1.64 were considered significant at *p* = .05 unilateral. Significant responses above noise level were recorded for every condition within the Left Parietal (minimal Z = 1.81), Right Parietal (minimal Z = 1.67), Medial Occipital (minimal Z = 5.22) and Medial Occipito-parietal regions (minimal Z = 2.08). We conducted a repeated-measures ANOVA on the Z values with condition (five levels), ratio (five levels), and electrode (four levels) as within-subject factors. The analysis showed a significant effect of electrode, *F*(3, 45) = 30.887, *p* < .001, confirming that the best responding electrodes were located within the Medial Occipital region. This posterior activity merely reflects the visual system’s synchronization to the stimulation. Neither the main effect of ratio nor any interaction did reach significance, all *Fs* < 1. Further, in the analysis, there was a significant effect of condition, *F*(4, 60) = 10.082, *p* < .001. However significantly different, brain responses were above the threshold value for every condition: mean Z for Area = 2.73, Convex Hull = 2.10, Number = 2.39, Size = 2.91, and Density = 2.12.

**Stimulus Assessment**

We conducted an independent assessment of images used in the conditions that had the largest ratio (1.5), by estimating the number of dots for each image, as well as four non-number dimensions: Convex Hull, Dot Size, Total Area, and Density. For each image, a mask was generated in which every non-background pixel was given the same value. Number of dots was then estimated as the number of contiguous regions in each mask, after first eroding the regions slightly to avoid counting abutting dots as a single region. The other parameters were similarly straightforward: The minimum number of pixels in the non-eroded contiguous region (“Dot Size”), the total number of non-background pixels (“Total Area”), convex hull of non-background pixels (“Convex Hull”), while “Density” was defined as convex hull divided by the number of dots (so in fact, inverse density). Together, these five factors provide a relatively complete description of the variability in the presented dot images.

**Table 1.** Average responses predicted by the Second Order Contrast model (SOC) in four primary visual areas. Brackets contain the upper and lower limits of 95% confidence intervals around the mean.

| **density** | *V1* | *V2* | *V3* | *V4* |
| --- | --- | --- | --- | --- |
| *standard* | 96.1 [82.4 114.0] | 97.9 [85.9 111.8] | 99.1 [86.7 111.9] | 98.6 [89.9 109.4] |
| *deviant* | 89.7 [93.1 95.5] | 92.1 [94.9 97.2] | 93.9 [95.9 98.6] | 96.7 [97.7 100.8] |
| **convex_hull** | *V1* | *V2* | *V3* | *V4* |
| *standard* | 99.7 [92.9 105.6] | 100.0 [94.6 103.7] | 99.7 [95.5 102.6] | 99.3 [97.0 101.4] |
| *deviant* | 103.0 [108.8 115.6] | 104.6 [109.4 114.4] | 106.3 [110.3 115.0] | 107.1 [109.6 113.1] |
| **area** | *V1* | *V2* | *V3* | *V4* |
| *standard* | 97.8 [90.0 106.9] | 98.1 [91.9 105.3] | 98.7 [92.8 105.2] | 98.6 [94.8 104.0] |
| *deviant* | 109.0 [113.6 114.7] | 105.7 [110.5 111.5] | 103.5 [108.2 109.4] | 101.6 [105.2 106.3] |
| **size** | *V1* | *V2* | *V3* | *V4* |
| *standard* | 100.9 [84.5 113.8] | 101.3 [86.3 112.9] | 101.6 [87.9 113.6] | 101.5 [91.5 109.8] |
| *deviant* | 95.8 [97.9 98.8] | 95.4 [97.3 98.9] | 95.3 [96.9 98.5] | 95.6 [96.8 98.5] |
| **number** | *V1* | *V2* | *V3* | *V4* |
| *standard* | 101.4 [87.5 107.0] | 100.9 [89.7 106.3] | 99.9 [91.6 105.5] | 100.4 [94.8 104.9] |
| *deviant* | 109.5 [111.0 112.4] | 106.3 [107.8 109.7] | 104.5 [106.4 108.3] | 102.3 [104.4 105.8] |
